# Multisite Evaluation of an Amplification-based Nanopore Sequencing Solution to Analyze Challenging Clinically Relevant Variants in Genes Associated with Hereditary Diseases

**DOI:** 10.64898/2026.05.14.725224

**Authors:** Stela Filipovic-Sadic, Connor A Parker, Mia K Mihailovic, John N Milligan, Jonathan M Turner, Stephanie L Borel, Vivian Le, Theodore Markulin, Justin W Janovsky, Bryan J Killinger, Malia J Deshotel, N. Scott Reading, Eric K Fredrickson, Yuan Ji, Devin Close, Jessica Wright, Mark Williams, Elizabeth S Barrie, Kendall E Martin, Shannon M Gray, Brian C Haynes, Bradley Hall

## Abstract

**Purpose:** Carrier screening for hereditary conditions is challenged by genes with complex genomic architecture, where short-read sequencing can fail to detect clinically relevant variants. This study evaluated a unified, amplification-based nanopore sequencing workflow across multiple laboratories for comprehensive analysis of such loci.

**Methods:** A modular long-read sequencing assay was evaluated across five laboratories using targeted PCR enrichment, Oxford Nanopore sequencing, and automated variant analysis. The workflow interrogated genes associated with spinal muscular atrophy, thalassemia, cystic fibrosis, fragile X syndrome, congenital adrenal hyperplasia, Gaucher disease, and hemophilia A. Performance was assessed against orthogonal methods for single nucleotide variants (SNVs), indels, copy-number variants, repeat expansions, and structural rearrangements.

**Results:** Across 882 unique samples (1,266 tests), overall agreement with comparator methods exceeded 96% for variant-level detection and 97% for genotype status classification. Long-read sequencing enabled phasing of paralogous loci, integrated sizing and interruption analysis for *FMR1* repeats, and simultaneous detection of SNVs and structural variants in globin loci and *CYP21A2*-*TNXB* region, reducing reliance on multiple workflows.

**Conclusion:** This multisite evaluation suggests that targeted long-read sequencing can consolidate complex variant detection into a single workflow, improving analytical completeness and operational efficiency for carrier screening.

## Introduction

Molecular genetic screening informs reproductive decision-making by identifying heterozygous carriers for autosomal recessive conditions and individuals with X-linked pathogenic variants.^1,2^ Next-generation sequencing (NGS) has broadened variant detection capabilities and is now widely used for genetic testing.^3^ However, conventional short-read platforms can underperform in regions with complex genomic architecture, such as those containing high homology, pseudogenes, repetitive elements, or structural variation.^4,5^ As a result, comprehensive analysis of these regions frequently requires multiple specialized, locus-specific assays. This fragmentation of testing workflows increases operational complexity, turnaround time, and analytical burden, while still risking incomplete detection of clinically relevant variants.^4,6^

Guidance from the American College of Medical Genetics and Genomics (ACMG) recommends an ethnicity-neutral, tiered approach to carrier screening, developed to support equitable access and clinical consistency.^1^ This framework considers factors including disease severity, carrier frequency, and clinical actionability and defines four screening tiers. Tier 3, currently encompassing approximately 97 autosomal recessive genes and 16 X-linked genes, is recommended for all individuals who are pregnant or planning a pregnancy.^1^ Within Tier 3, ACMG laboratory technical standards further identify a subset of genes as technically challenging due to complex genomic architecture.^4^ Variant detection in these regions remains challenging for short-read sequencing, as alignment ambiguity can obscure variant origin and phasing, often necessitating ancillary methods to achieve desired clinical sensitivity.^4,6^

Recent advances in long-read sequencing technologies, particularly those from Oxford Nanopore Technologies (ONT) and Pacific Biosciences (PacBio), have begun to address long-standing gaps in complex variant detection with short-read sequencing. These single-molecule systems generate reads that can exceed tens of kilobases, enabling simultaneous detection of SNVs, indels, short tandem repeats (STRs), copy number variants (CNV), and structural variants (SVs) within a unified workflow. By spanning repetitive and homologous regions, long reads can facilitate direct phasing, pseudogene resolution, accurate characterization of large SVs and STRs, thus reducing the need for reflex testing.^7–12^ By integrating enrichment, per-sample barcoding, sequencing, and downstream analysis, these approaches can support multiple samples within a modular and scalable framework.^13–15^

The genes evaluated in this study were selected based on both technical complexity and relevance to commonly implemented ACMG-compliant carrier screening workflows, collectively representing a broad spectrum of clinically relevant variant types. These include paralogous genes subject to copy number variation, such as *SMN1* (HGNC:11117) and *SMN2* (HGNC:11118), where accurate copy discrimination is critical for assessment of spinal muscular atrophy (OMIM:253300).^13,16^ Loci with highly homologous pseudogenes, including *GBA1* (HGNC:4177), associated with Gaucher disease (OMIM:230800), and *CYP21A2* (HGNC:2600), associated with congenital adrenal hyperplasia due to 21-hydroxylase deficiency (OMIM:201910), present additional analytical challenges arising from frequent gene–pseudogene conversions and complex rearrangements.^17–19^ The *CYP21A2* locus is further complicated by its genomic proximity to *TNXB* (HGNC:11976), where recombination events can generate chimeric alleles associated with CAH-X syndrome (OMIM:617056).^12^ Structural variation and copy-number changes within the α- and β-globin gene clusters (*HBA1*, HGNC:4823; *HBA2*, HGNC:4824; *HBB*, HGNC:4827) underlie α- and β-thalassemia (OMIM:604131 and OMIM:613985, respectively) and often require multiple complementary assays for resolution.^20^ Repeat expansion loci such as *FMR1* (HGNC:3775), associated with fragile X syndrome (OMIM:300624), require precise sizing of CGG repeats and phasing with AGG interruptions to inform expansion risk.^21–23^ Finally, large intronic inversions in *F8* (HGNC:3546), a common cause of hemophilia A (OMIM:306700), are typically undetectable by short-read sequencing and rely on inversion-specific methods.^24,25^

To address these challenges, we implemented a modular, kit-based workflow that combines short- and long-range PCR enrichment with nanopore sequencing and evaluated it across 5 laboratories. This approach enables streamlined genotyping of genes typically well resolved by short-read sequencing (*CFTR* and *HBB*) alongside analytically challenging loci (*SMN1/SMN2*, *FMR1*, *HBA1/HBA2*, *GBA1*, *CYP21A2*, *TNXB*, and *F8* intronic inversions). Collectively, these targets account for approximately 70% of at-risk couples identified through carrier screening.^26^ By comparing locus-specific performance to established orthogonal methods, this multisite evaluation assesses the feasibility of consolidating technically challenging variant detection within a single workflow, with the potential to reduce reliance on fragmented, assay-specific testing approaches.

## Materials and Methods

### Study Design and Assay Workflow

This study evaluated a standardized, amplification-based long-read sequencing workflow across five laboratories, including one developer site and four independent external sites. All sites followed a common protocol and used the same assay reagents and automated analysis software, while testing was performed on site-specific but comparable laboratory instrumentation.

The assay employs a modular design consisting of four reagent kits that can be used individually or in combination: Kit A (*CFTR, SMN1, SMN2*), Kit B (*FMR1*), Kit C (*HBA1, HBA2, HBB*), and Kit D (*CYP21A2, TNXB, F8* inversions, *GBA1*), enabling interrogation of multiple variant classes within a unified workflow (Figure 1A). The workflow comprises three main phases: targeted short- and long-range PCR enrichment, nanopore sequencing, and automated variant analysis (Figure 1B). A related workflow applied to *SMN1/SMN2* analysis has been previously described.^13^ This study extends that framework to additional clinically relevant loci, implements automated variant analysis, and evaluates performance across five independent laboratories.

**Figure 1.**
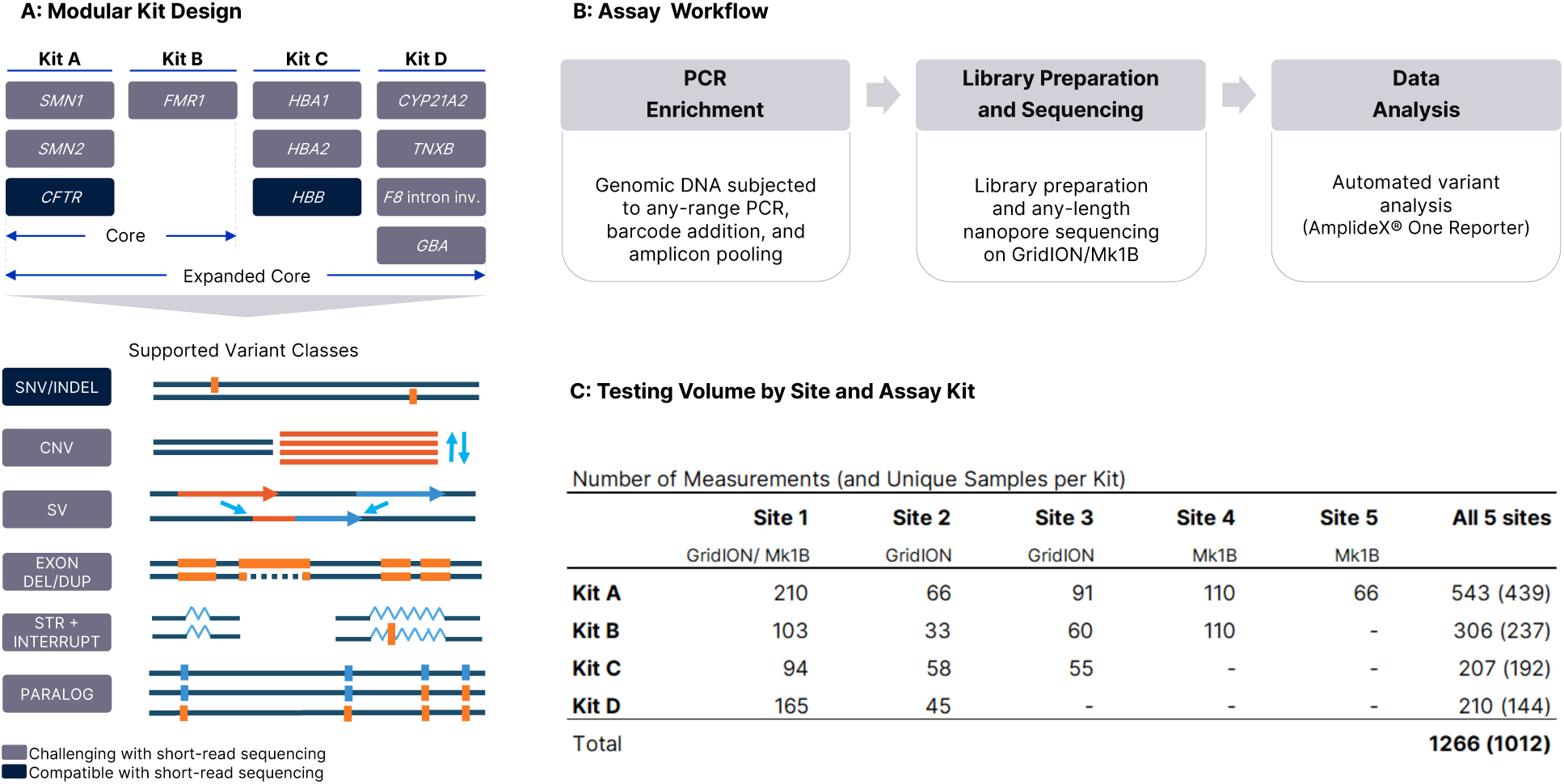
Assay design, workflow, and study scale. **(A)** Modular kit design showing gene targets included in Kits A–D. Kits can be used individually or in combination to genotype each sample. In multi-gene kits, analysis module settings allow reporting to be restricted to a single gene if desired. Kits A (*CFTR, SMN1, SMN2*) and B (*FMR1*) target core genes widely implemented in clinical carrier screening (*CFTR* and *SMN1* are recommended as Tier 1 in the ACMG 2021 carrier screening recommendations; *SMN2* is included for modifier copy assessment; *FMR1* is categorized as Tier 3 but is broadly included in core carrier screening panels). Kit C (*HBA1, HBA2, HBB*) and Kit D (*CYP21A2, TNXB, F8* intron inversions, *GBA1*) expand coverage to additional clinically important loci characterized by complex genomic architecture and diverse variant classes. Genes and variant classes are color-coded to indicate analytical complexity. Variant classes detected by the unified workflow include single-nucleotide variants (SNV), small insertions/deletions (INDEL), copy-number variants (CNV), large structural variants (SV), exon-level deletions/duplications (exon del/dup), short tandem repeats (STR) with interruptions, and variants phased to paralogous or pseudogene-associated loci (PARALOG). **(B)** Unified assay workflow: Genomic DNA was subjected to long-range multiplex PCR enrichment, barcoding, pooling, and library preparation using ONT ligation chemistry, followed by sequencing on GridION or MinION Mk1B instruments. Primary analysis (basecalling and demultiplexing) was performed in MinKNOW; secondary analysis (alignment, read depth and other quality controls, variant identification, variant classification, data visualizations, and reporting) was performed using AmplideX One Reporter with integrated Nanopore Carrier Plus Analysis Module. **(C)** Summary of testing volume across five sites, stratified by kit. Values represent total measurements, with unique sample counts in parentheses. Total of 882 unique genomic DNA samples were utilized in this study, with select samples analyzed in multiple kits, or across sites, resulting in 1012 unique sample-kit pairs. A subset of samples was also tested in replicate at individual sites, resulting in 1266 total measurements.

Targeted PCR enrichment, generating amplicons approximately 0.3 to 10 kb in length, was performed using the AmplideX® Nanopore Carrier Plus Kits A-D according to the manufacturer’s instructions (Asuragen, a Bio-Techne brand; Austin, TX, USA). The accompanying kit protocol specifies all required wet-lab processing steps and materials throughout the workflow, including library preparation and sequencing. In brief, following gene-specific amplification, samples were PCR-barcoded (PCR Barcode 96 Plate; ONT, Oxford, UK), pooled, and converted to sequencing libraries using ligation chemistry (Ligation Sequencing Kit SQK-LSK114; ONT). Libraries were sequenced on R10.4.1 flow cells (FLO-MIN114) using MinION Mk1B or GridION instruments (ONT).

Basecalling and barcode demultiplexing were performed in MinKNOW using Dorado basecaller (ONT, DNA SUP v4.3 model). FASTQ files were analyzed using AmplideX^®^ One Reporter v1.0.10 with Nanopore Carrier Plus Analysis Module v1.0.4, which automated alignment to the GRCh38 human reference genome, allele deconvolution, variant calling, and quality control. Automated outputs included VCF files, tabulated summaries at the sample, gene, and variant level, and graphical reports for laboratory review.

### Samples and DNA Input

Archived residual clinical samples and cell line-derived DNA were used in this study. All samples were de-identified, and no patient identifiers were available to investigators involved in centralized data aggregation or analysis. Genomic DNA was isolated from peripheral blood using validated extraction methods, including salting-out/precipitation, silica-column, and magnetic-bead–based chemistries. Cell line-derived DNA was obtained either from Asuragen (Austin, TX, USA) or the NIGMS Human Genetic Cell Repository at the Coriell Institute for Medical Research (Camden, NJ, USA). DNA purity and yield were assessed using a NanoDrop™ spectrophotometer (Thermo Fisher Scientific, Waltham, MA, USA). Each reaction was prepared using a target DNA input of 40 ng, with an acceptable range of 40–100 ng.

A total of 882 unique genomic DNA samples were interrogated across five laboratories, including 150 cell line-derived and 732 whole blood-derived samples, excluding calibrators and optional no-template controls. A subset of samples was tested at more than one site or in more than one kit, resulting in 1012 unique sample-kit pairs across sites. In addition, some sites analyzed select samples in replicate, resulting in 1,266 total measurements across the four assay kits (Figure 1C).

Orthogonal data were obtained using methods routinely employed for each gene at the participating sites. These methods varied by gene, variant class, and laboratory, encompassing 49 assays across 16 method types, reflecting the diversity and complexity of current clinical testing practices (Table S1).

### Bioinformatics Pipeline

MiniMap2^27^-derived long-read alignments were tagged and filtered for amplicon assignment before routing to individual analysis modules (e.g., SNV/indel, copy number (CN)). For loci in which copy number calls greater than 2 were supported (*SMN1/2* in Mix A, *HBA* in Mix C, and *CYP21A2, GBA1*, and *TNXB* in Mix D) or for repeat-length estimation (*FMR1*), the analysis module separated amplicon-specific reads into allele groups prior to variant detection, either through sequence deconvolution or length binning, respectively. Single nucleotide variants and indels were called using Clair3^28^ on either combined reads or allele-specific read groups, as applicable. Larger structural variants were analyzed with Sniffles2.^29^ Custom models and algorithms were developed to call copy number variants, repeat expansions (e.g., *FMR1*), and gene conversions or deletions in loci with high sequence homology to pseudogenes (e.g., *SMN1/SMN2, CYP21A2*). Results were annotated with ClinVar^30^ data accessed on March 21, 2025, subjected to in-silico variant effect prediction (VEP)^31^, and visualized with integrated coverage and variant plots.

### Data Analysis

QC-passing variant calls were compiled from the five sites and genotype calling performance was summarized on a per-site and collective basis. For samples with genotypes reported by orthogonal assays, variant level performance was evaluated for the following genes and genotype categories: STR (*CFTR, FMR1*), CNV (*CFTR, HBA1/HBA2, HBB, SMN1, SMN2*), SNV/INDEL (*CFTR, SMN1/2, HBA1/2, HBB, CYP21A2, GBA1*), and inversions (*F8*), using variant-appropriate units of comparison.^32^ Orthogonal data for *TNXB* exon 32-44 was not available, therefore performance was excluded from this study.

For SNV/indel analysis, detection sensitivity and specificity were compiled on a per-variant, per-allele basis (regardless of annotated pathogenicity).^33^ Heterozygous variants were evaluated with flexible thresholds accounting for gene-specific copy number, in which any number of variant-containing copies were considered true positive (TP) if the total variant copies detected was fewer than the total number of amplicon copies detected. *SMN1/2* SNV/indel results were analyzed independently for Mix A and Mix D whenever both were utilized for the same sample, with Mix D providing longer amplicons that support more comprehensive phasing. Accurate phasing of compound variants was not evaluated in this analysis but was incorporated into downstream carrier genotype status interpretation (Figure 2). For *FMR1*, CGG repeat categories were assigned according to ACMG guidelines^23,34^ based on the size binning of the largest detected allele. CGG-repeat length and AGG-interruption position were evaluated for accuracy within size precision (Table S2) relative to the closest expected allele lengths and allele-specific interruption positions, respectively. AGG calls from full-mutation alleles were excluded from performance analysis due to lack of both orthogonal data and clinical utility of risk.^21^

**Figure 2.**
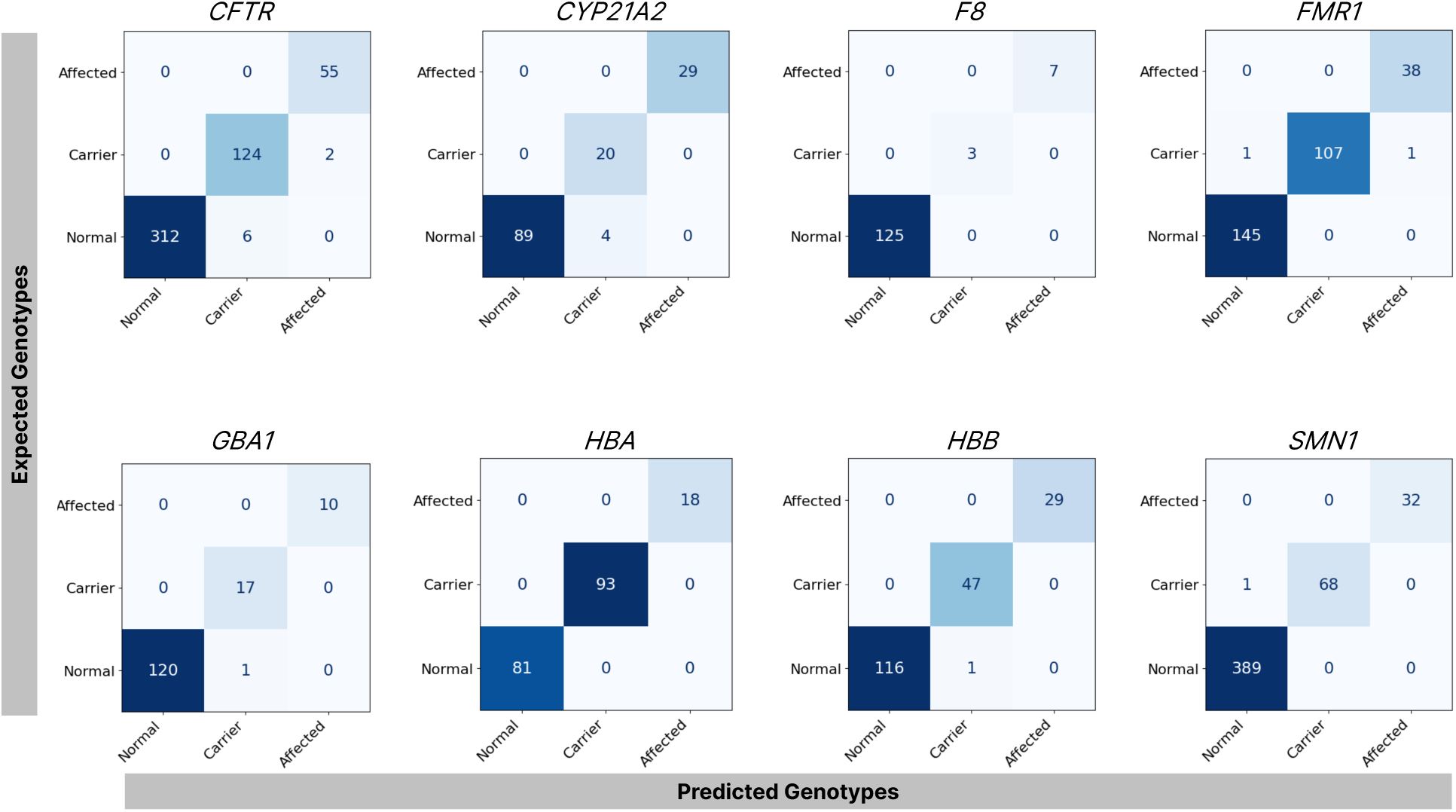
Genotype status agreement between the assay and orthogonal methods. Confusion matrices show the number of samples classified as Normal, Carrier, or Affected by both methods for each gene, with rows representing orthogonal method classifications and columns representing assay classifications. Overall percent agreement (OPA) was calculated per gene within each assay panel: Mix A (*CFTR*- 98.4% OPA, *SMN1*-99.8% OPA), Mix B (*FMR1*-99.3% OPA), Mix C (*HBA1/2*- 100% OPA, *HBB*- 99.5% OPA), and Mix D (*CYP21A2*- 97.2% OPA, *F8*-100% OPA, *GBA1*- 99.3% OPA).

In addition to variant-level performance, assay results were evaluated at the level of predicted genotype status, reflecting the categorical interpretation typically reported in clinical carrier screening. For each gene, genotype outputs were transformed into categorical outcomes of Normal (NOR), Carrier (CARR), or Affected (AFF). The pipeline results were generated using the default analysis filter, which only summarizes SNV/-indel whose ClinVar^30^ annotations are categorized within the software as pathogenic or likely pathogenic (P/LP) according to ACMG guidelines.^33^ In contrast to variant level analysis, samples were conservatively presumed normal for each gene that lacked gene-specific orthogonal data. For cases in which genotype status is determined by gene copy number (*CYP21A2*, and *GBA1)*, these categories were assigned based on the number of P/LP variant-free gene copies detected (NOR = ≥2; CARR = 1; AFF = 0). For *HBA*, genotype status category was similarly assigned on the basis of *HBA1*+*HBA2* P/LP variant-free copies (NOR = ≥4; CARR = 2-3; AFF = ≤1). For *CFTR, F8, FMR1, HBB*, and *SMN1*, genotype categories were assigned using gene-specific interpretation rules applied to the genotype summary column as outlined in Table S3. Special consideration was made for the following cases: (1) for multi-amplicon gene targets (*CFTR, HBB*), the presence of multiple, amplicon-unique heterozygous P/LP SNV/indels was assumed to be in trans and categorized as AFF, (2) Genotype status of samples with a predicted *F8* inversion of ambiguous genotype (i.e., Unknown as opposed to Homozygous/Heterozygous) were assigned Carrier or Affected on the basis of sex, (3) *CYP21A2* copy number truth was excluded from this transformation due to the abundance of discordant variants that appeared derived from *CYP21A1P* in information-limited NGS-based orthogonal assays that cannot phase the entire gene. Discrepant results without orthogonal confirmation were excluded from the main figures (Figure 2) but included in Figure S1 for transparency. This exclusion was implemented to avoid bias introduced by NOR assumptions in cases lacking complete comparator data.

## Results

To evaluate overall assay performance, we assessed cross-site concordance at the variant and genotype-status levels, followed by qualitative analyses highlighting locus-specific insights uniquely enabled by long-read sequencing.

### Variant Level Agreement

Variant-level agreement with available orthogonal data was assessed per gene and variant type across all laboratory sites and within each site (Table S4). Assay performance was consistent across internal and external sites, with overall agreement 96.12% to 100% across genes and variant types. (Table 1).

**Table 1.**
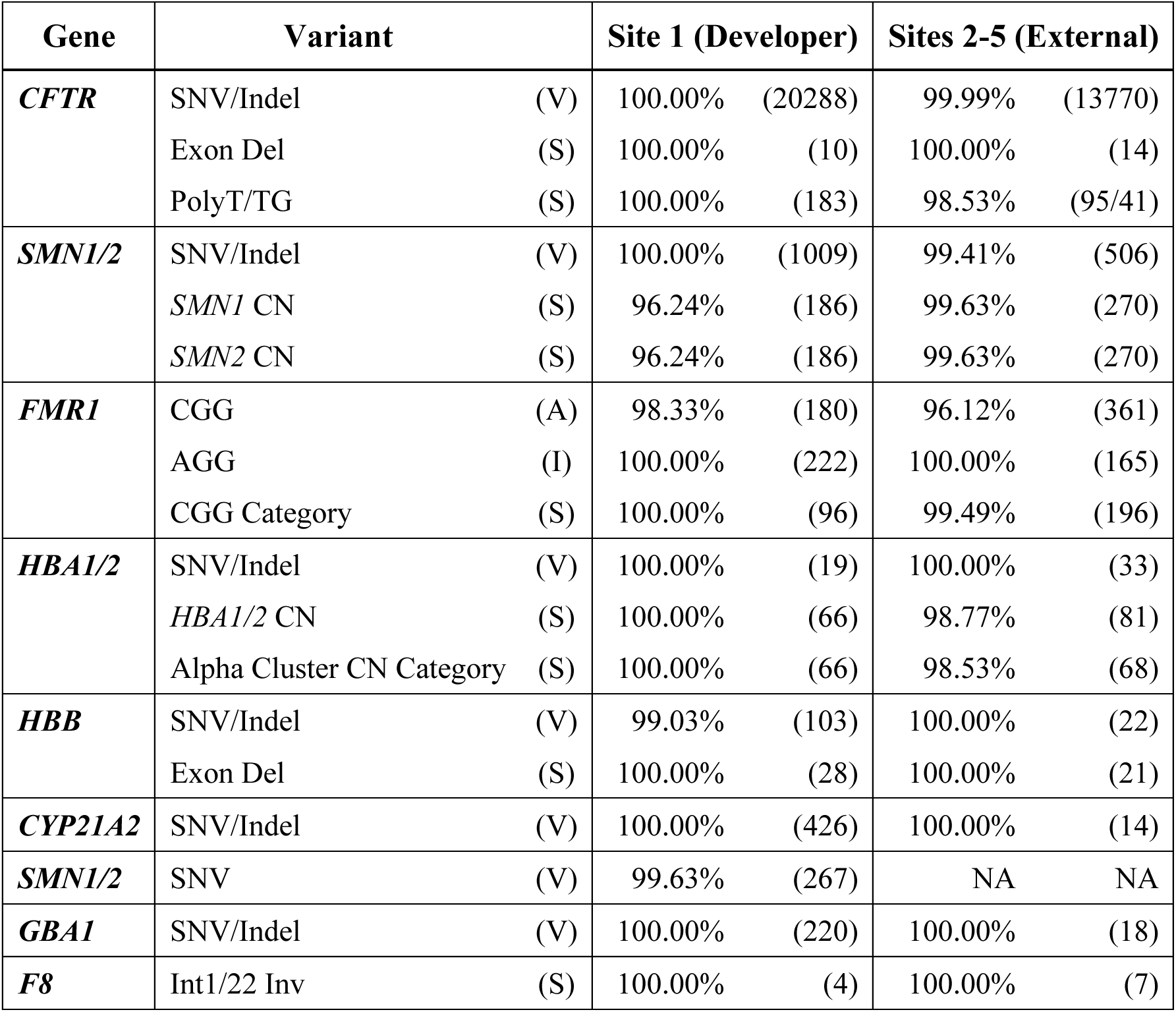
Variant-level agreement by gene and variant class across developer and external laboratories. Overall percent agreement (OPA) between the assay and orthogonal methods is shown for each gene and variant class across five laboratories. Site 1 represents the developer laboratory, and Sites 2–5 represent aggregate external laboratories. Columns show OPA with orthogonal methods, followed by the number of measurements in parentheses. Variant types include SNVs/indels, copy numbers (CNs), large structural variants (SV), short tandem repeats (*FMR1* CGG and *CFTR* poly-T/TG) and interruptions (AGG), and category-level classifications (e.g., CGG category, alpha cluster CN category). OPA is equivalent to positive performance agreement (PPA) for select analyses (*i.e., CFTR* exon deletions and *F8* intron 1 and 22 inversions) due to the absence of orthogonal confirmation of negative samples. Abbreviations: V = variant-level concordance; S = sample-level concordance; A = allele-level concordance; I = interruption-level concordance; Exon Del = large exon deletion; PolyT/TG = poly-T/TG tract; Int1/22 Inv = intron 1/intron 22 inversion. NA = data not available or not applicable for a given site or variant type.

To better understand the underlying causes of the few observed discrepancies, we performed a detailed review of discordant calls across genes and variant types. Variation in *SMN1* and *SMN2* copy number agreement (96.24–100%) was mainly attributable to discrepancies in detecting ≥3 copies versus 2 copies. Across all sites, only a single discrepancy was observed out of 456 measurements at clinically relevant *SMN1* copy number levels of 0 and 1, in which a 1 copy sample was reported as 2 copies. All SNV/Indel discrepancies for *SMN1* were attributed to limitations of the orthogonal method to accurately phase calls with the correct paralog. Specifically, the assay phased a small 3′ untranslated region deletion to the *SMN2* paralog in multiple replicates of two different samples, whereas the orthogonal assay reported the variant on *SMN1* for both samples, which can indicate a 2 + 0 genotype in certain ethnic groups as previously described.^13,35^

Overall, SNV/Indel detection was robust with >99% agreement for each gene at internal and external sites. A single *CFTR* SNV was missed by the assay due to suspected partial gene duplication affecting the *CFTR* region that creates an imbalanced allelic ratio that confounds standard diploid variant calling assumptions. For *FMR1* CGG repeat detection, a slightly higher agreement was observed internally (98.33% vs 96.12%), which can be explained by annotation and subsequent reconciliation of low-frequency alleles (due to mosaicism), which was not performed at external sites due to orthogonal assay data limitations. For *HBB*, a heterozygous SNV detected by Sanger sequencing was discordant with a hemizygous call by the assay in a beta thalassemia clinical sample. This variant was observed in parallel with deletion of the exon that contains the variant loci. We hypothesize that the detected exon deletion is partial and does not encompass the region amplified in Sanger sequencing allowing detection of both variant and wild-type alleles by Sanger sequencing.

### Genotype Result Agreement

Gene-level genotype status accuracy was determined to evaluate the expanded phasing capabilities offered by the assay’s incorporation of long-read sequencing. This analysis extended the performance assessment to a larger dataset than variant level analysis by incorporating presumed normal genotypes for samples where orthogonal data was unavailable. Overall agreement between methods was >97% across the eight gene categories assessed, with concordance ranging from 97.2% (*CYP21A2*) to 100% (*HBA1/2, F8 intron inversion*; Figure 2).

Across all genes and sites, only two false negative (FN) calls were observed. One FN occurred in *FMR1* and was attributable to low-level size mosaicism in which the orthogonal method detected a low-frequency 56-repeat premutation allele changing categorical bounds into the premutation category, whereas the nanopore assay detected only the two predominant alleles (29 and 48 repeats, each with two AGG interruptions). The 56-repeat premutation allele was visible in the sequencing repeat size read depth histogram reported by the software but was below the call threshold. The second FN occurred in *SMN1*, where the assay reported two gene copies for a sample in which orthogonal results indicated a single copy, as mentioned in variant level accuracy.

False positives (FP) were limited to large deletions detected in *CFTR*, *HBB*, *GBA1*, and *CYP21A2* and a single full mutation in *FMR1*. For *CFTR*, seven of the eight FP calls were exon 19-20 deletions and the other was a full gene deletion. Automatic reporting of large exon deletions by the assay within *CFTR* requires detection in at least 2 consecutive amplicons, except for the single amplicon covering both exon 19 and 20. A single FP for an *HBB* exon 3 deletion was caused by an alpha cluster duplication and increased signal from the normalization region used in the detection algorithm. A single FP for *GBA1* and four FP for *CYP21A2* full gene deletions were in samples with likely homologous *GBA1* and *CYP21A2* alleles. Sequence deconvolution of gene sequence (*CYP21A2* or *GBA1)* into allele groups and subsequent allele group read depth normalization to the pseudogene (*CYP21A1P* or *GBAP*) is utilized to determine the copy number of *CYP21A2* and *GBA1,* which can be confounded when alleles are homologous and/or pseudogene copies are absent or homologous. In *FMR1*, a single sample was reported with a FP full mutation call in addition to accurate detection of a known premutation allele (149 CGG). It is unclear whether this discordance reflects sensitive detection of low-level size mosaicism or arises from low-level noise in read depth.

Collectively, these findings demonstrate high cross-gene concordance and consistent genotype status determination across laboratories. The discrepancies were primarily traceable to large structural variants or rare mosaic configurations, which may be improved by assay and analysis refinements in future versions.

### Greater Insights

In addition to strong variant and genotype-level performance, data for several genes provided additional qualitative insights beyond the capabilities of traditional methods. Specifically, data from loci including *HBA1/2* (Figure 3), *CYP21A2* (Figure 4) and *FMR1* (Figure S2), demonstrated how structural resolution, graphical visualization, and direct long-read phasing can enhance variant interpretation while reducing reliance on multiple assay workflows.

**Figure 3.**
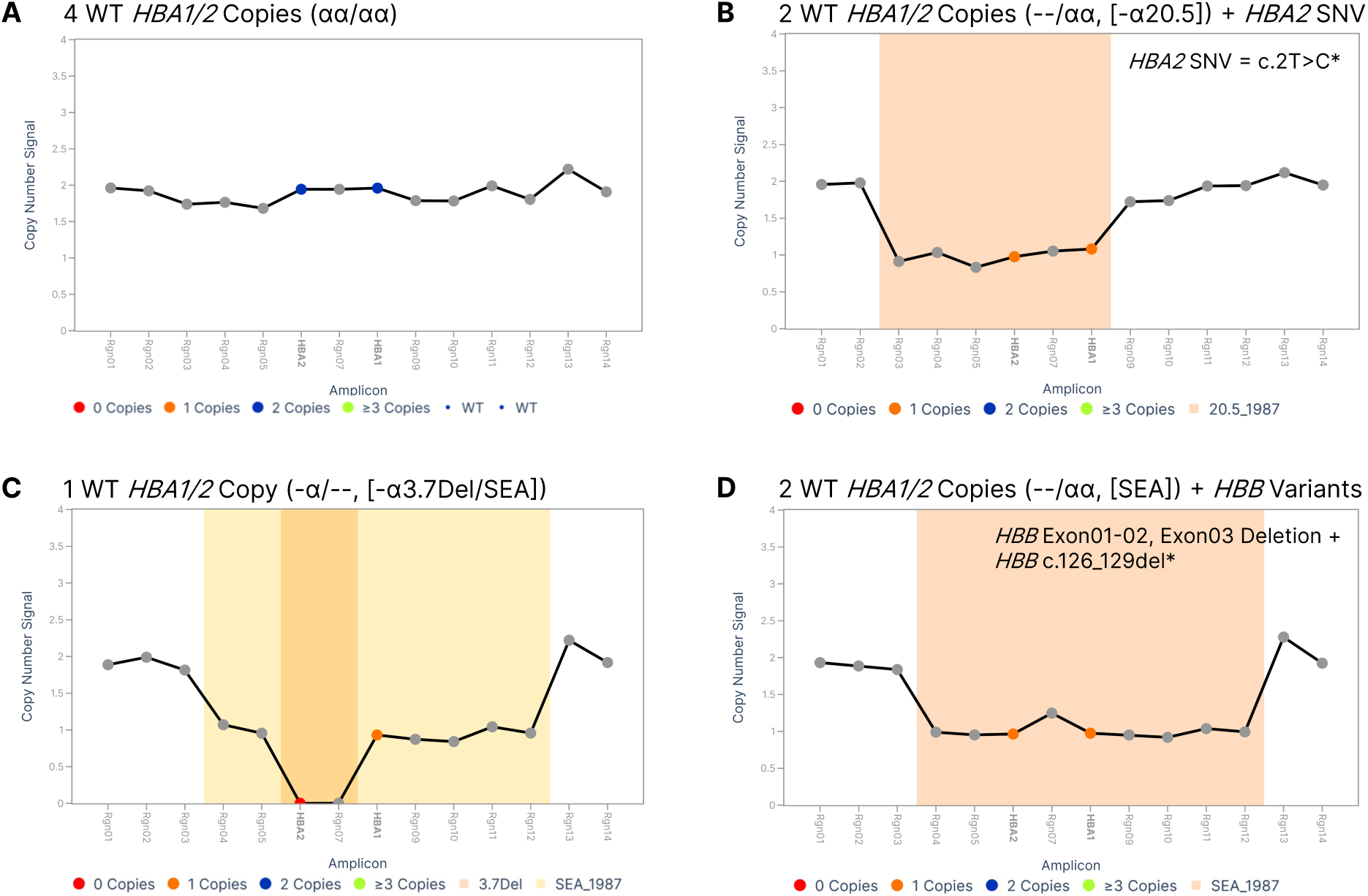
Integrated copy-number and sequence variant resolution across the α-globin cluster using long read nanopore sequencing. Long read sequencing of the complete *HBA1* and *HBA2* genes, combined with targeted amplification across the α-globin cluster, enabled simultaneous detection of large structural variants and *HBA1/2* SNV/INDELS. **(A)** Normal sample with four *HBA1/2* copies (αα/αα). **(B)** Concurrent detection of copy number changes and pathogenic variants, illustrated by a sample with two wildtype *HBA1/2* copies due to a 20.5 kb deletion and an *HBA2* SNV. **(C)** identification of multiple structural variants (3.7 kb deletion and SEA deletion) within a single sample, and **(D)** concurrent assessment of *HBB* gene status (*HBA1/2* SEA deletion + *HBB* gene deletion/*HBB* INDEL, *HBB* data not shown). Deletion nomenclature includes the year of first description where applicable (e.g., 20.5_1987, SEA_1987). CN = copy number; SNV = single-nucleotide variant; INDEL = insertion/deletion. *SNV/INDEL data not shown.

**Figure 4.**
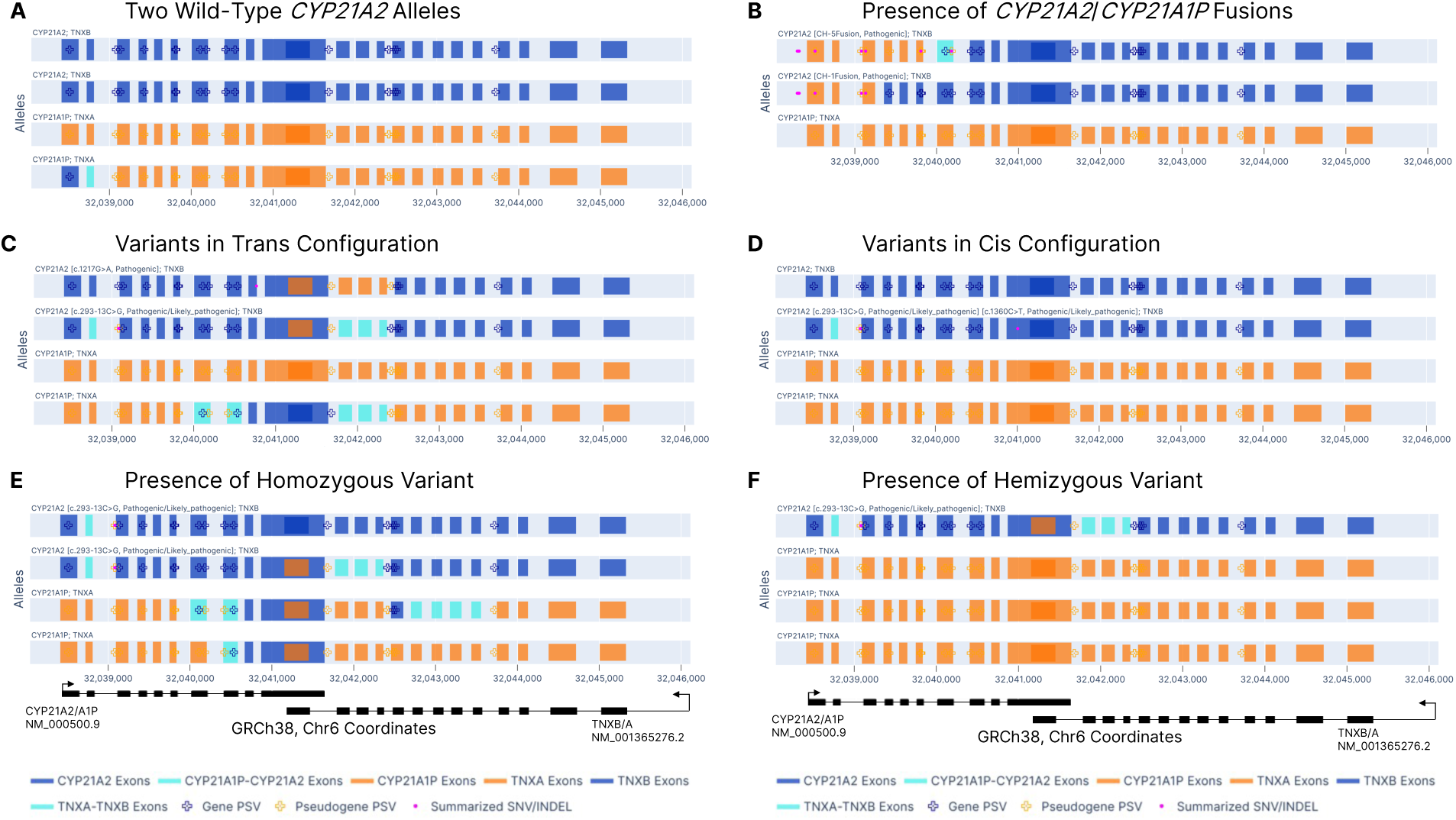
Integrated detection of *CYP21A2* sequence, structural, and copy-number variants. Primers amplify both gene/pseudogene of the full *CYP21A2* gene and exons 33–44 of *TNXB and CYP21A2/TNXA* to enable allele-level grouping by alignment and sequence deconvolution and variant analysis. Grouped alleles are graphically represented: **(A)** differentiation of *CYP21A2* from its pseudogene *CYP21A1P* for a wild-type sample, **(B)** identification and classification of *CYP21A2*/*CYP21A1P* chimera alleles (CH-1 through CH-7, CH-9) resulting from large deletions, **(C–D)** differentiation of compound heterozygotes from heterozygotes via phasing of variants to distinguish trans or cis configurations, and **(E–F)** discrimination between homozygous and hemizygous variants due to copy number differences. Blue (gene exons) and orange (corresponding pseudogene regions) are differentiated by alignment and known PSVs (represented by plus icon) whereas undifferentiable regions (teal) have no or conflicting PSV origin within the exon region. SNVs (pink dots) are positioned in the graph on alleles where sequence aligns to the gene. Genomic coordinates are listed for the *CYP21A2/TNXB* genomic region only. Arrows indicate *CYP21A2* and *TNXB* coding direction. PSV = paralogous sequence variant; SNV = single-nucleotide variant; INDEL = insertion/deletion.

#### HBA1/2 - simultaneous α- and β-globin cluster characterization

Molecular evaluation of thalassemia often requires a cascade of tests that include copy number evaluation (typically by multiplex ligation-dependent probe amplification or gap-PCR) and SNV analysis (typically by Sanger sequencing or NGS) to resolve genotype of *HBA1, HBA2*, and *HBB*, which encode the α- and β-globin subunits of hemoglobin.^36^ Recently, a long-read sequencing prototype has been proposed as a comprehensive method, but with limitations in differentiating the seven most common SVs.^20^ The design and workflow described here enabled simultaneous interrogation of SNVs and SVs in *HBA1, HBA2*, and *HBB*, including differentiation of the common alpha globin structural variants within a single assay.

Representative examples include visual representations of a wild-type sample (Figure 3A) and three samples with structural variants leading to copy number variations across the α-globin cluster (Figure 3B-D), allowing comprehensive characterization of genetic alterations associated with α- and β-thalassemia. For example, in Figure 3B, a sample having a single copy of *HBA1* and *HBA2* was correctly identified with the –(α)^20.5^ deletion. In addition, the remaining *HBA2* copy was shown to harbor a rare pathogenic SNV, indicating that the sample contained only a single functional alpha-globin gene copy. A second sample with a single functional alpha-globin gene copy (Figure 3C) was correctly identified via simultaneous detection of two structural variants (3.7 kb deletion and SEA deletion). Another sample demonstrated the assay’s ability to resolve complex multi-gene alterations: the sample was correctly identified with a loss of one *HBB* gene copy and the remaining *HBB* allele contained a pathogenic small indel, resulting in no functional *HBB* copies (Figure 3D). The same sample also exhibited a single copy of *HBA1* and *HBA2* due to the presence of the SEA deletion, which has been associated with reduced beta-thalassemia severity due to increased balance of the alpha/beta subunits.^36^ Besides detection and classification of widely annotated α-globin cluster SVs, the assay detected an *HBA1/2* SVs outside this set, with approximate breakpoints inferred from assay outputs (Figure S3).

#### *CYP21A2* - Integrated detection and phasing of SNVs, CNs, and chimeric alleles

The *CYP21A2/TNXB* locus is part of the bi-modular RCCX domains 30 kb apart from the pseudogenes *CYP21A1P/TNXA* that share ∼98% sequence homology. Pathogenic variants result from rearrangements due to unequal recombination, such as microconversions resulting in SNVs or larger mono/tri-modular SVs resulting in copy number changes or chimeric alleles.^38^ Conventional approaches often require multiple assays, including long-range PCR followed by Sanger sequencing for SNVs, MLPA for copy number changes, and targeted PCR to detect gene conversions for accurate interrogation of this genetically complex locus.^37,38^

In this study, long-read sequencing combined with automated sequence deconvolution enabled simultaneous detection and visualization of wild-type samples (Figure 4A) and various pathogenic SNV/indels, gene copy number, pseudogene content, and structural rearrangements (Figure 4B-F), consistent with recent long-read studies of *CYP21A2*.^39,40^ For example, a sample containing two *CYP21A2/CYP21A1P* chimera alleles and zero wild-type *CYP21A2* alleles was accurately identified and annotated according to known chimera sub-types (CH-1 and CH-5) (Figure 4B). In addition, phasing of two pathogenic variants in trans or cis configurations across reads >10kb was also achieved (Figures 4C and 4D respectively). Concomitant sequence analysis and copy number detection distinguished between two samples with homozygous versus hemizygous variants in which all *CYP21A2* alleles contain the same pathogenic variant but different copy numbers of *CYP21A2* (Figure 4E and 4F respectively). Similar allele-level resolution has been reported in recent long-read CAH cohorts to inform genotype interpretation at this complex locus.^39,40^ Collectively, these examples demonstrate enhanced variant detection across this complex genomic region relative to conventional assays.

## Discussion

This multisite evaluation demonstrates that a cohesive targeted PCR-enrichment nanopore sequencing assay can serve as a comprehensive platform for unified detection of diverse variant classes including SNVs/indels, copy number changes, large deletions, repeat expansions with interruptions, gene conversions, and structural inversions with a single workflow. The approach addresses the central operational challenge in this space: complex loci historically require fragmented workflows that increase cost, prolong turnaround time, and create additional opportunities for analytical or interpretive errors. The results across five laboratories demonstrate that performance rivals current fragmented methods, with >96% agreement at the variant level and >97% concordance for genotype calls.

The workflow presented herein incorporates widely accessible nanopore long-read sequencing platforms including MinION Mk1B and GridION devices, and automates analysis from basecalling, through variant classification and genotype reporting to reduce bioinformatics burden. The data outputs include standard VCF formats, tabular reports, and supporting figures for simple genotype interpretation that enables LIMS integration and tertiary classification or reporting.

This study demonstrated modularity and scalability for samples and genes interrogated. The standardized enrichment chemistry, implemented as four independent mixes, was combined by different sites in various combinations for single sequencing runs. Furthermore, varying sample numbers (Table 1) were enriched in single sequencing batches between 10 and 96 samples. This adaptability allowed external sites to align sequencing capacity with workflow volume or resource constraints.

Several opportunities for further refinement were identified. To support high-throughput screening (beyond 96 samples per run), enhancements to software and workflow, including dual barcoding and PromethION compatibility, could be incorporated. Additional software updates to enable automated detection of specific variants are planned. For instance, fusions implicated in CAH-X, a condition with characteristics of congenital adrenal hyperplasia and Ehlers-Danlos syndrome can currently only be manually detected. Although the assay produces sufficient read depth in a single amplicon across the *CYP21A2* gene and exons 32–44 of *TNXB* to resolve these fusions, the present study included only a limited number of samples with established truth for *TNXB* and CAH-X genotypes (n=4). Larger studies incorporating well-characterized samples and additional positive cases may be required for comprehensively evaluating performance in this region and refining algorithmic detection strategies. Another potential improvement includes CN and structural variant modeling. Occasional false positives and normalization artifacts observed in large exon deletion calls (e.g., single exon deletions in *CFTR*) highlight the need for refined CN models trained on representative data sets. Similarly, algorithmic improvements for detecting duplications and deletions within the α-globin cluster may be necessary to enhance copy number accuracy for *HBA1/2* and *HBB*, where structural changes can occasionally affect normalization regions used in CN analysis.

As long-read sequencing technologies continue to mature, targeted workflows such as this one represent a pragmatic bridge between analytical completeness of whole genome long-read sequencing and operational feasibility of current workflows in high and low volume laboratories. Continued optimization of enrichment chemistries, variant-calling algorithms, and content expansion may enable broader inclusion of technically challenging genes within routine clinical testing.

In conclusion, this study establishes the first modular, amplification-based long-read nanopore sequencing panel within a unified workflow to enable accurate detection of diverse variant types across multiple laboratories. By integrating automated complex variant analysis into the assay, the approach addresses key unmet needs in carrier testing traditionally achieved through fragmented workflows and supports the expanding role of long-read sequencing in clinical genomics.

## Supporting information

Coriell Cell Line Data

## Data Availability

The data that support the findings of this study are available from the corresponding author upon request.

## Acknowledgements

We would like to thank Alina Ham, Sandra Guera-Roberts and Rebecca Smith from Oxford Nanopore Technologies, and Jacob Ashton, Elliot Hallmark and Ninad Pendse from Asuragen for their collaboration and support during this study.

## Author Contributions

Conceptualization: B.H., B.C.H., J.N.M, S.F.S., C.A.P, M.K.M.; Data curation: C.A.P.,M.K.K, J.M.T, S.L.B., V.L., T.M., J.W.J., B.J.K., M.J.D., J.W.J., K.E.M., S.M.G.; Formal analysis: C.A.P, M.K.M.; Investigation: J.M.T, S.L.B., V.L., T.M., J.W.J., B.J.K., M.J.D., N.S.R., E.K.F., Y.J., D.C., J.W., M.W., E.S.B., K.E.M., S.M.G., B.C.H., B.H.; Methodology: S.F.S., C.A.P, J.N.M, J.W.J.; Project Administration: S.F.S.; Resources: S.F.S., B.J.K., D.C., M.W., E.S.B., S.M.G., B.C.H., B.H; Software: T.M., B.J.K.; Supervision: J.N.M, B.J.K., D.C., M.W., E.S.B., S.M.G., B.C.H., B.H.; Visualization: S.F.S., C.A.P, M.K.M, J.N.M, B.C.H., B.H.: Writing-original draft: S.F.S., C.A.P, M.K.M, B.H., J.N.M; Writing-review & editing: all authors.

## Ethics Declaration

Use of archived, de-identified residual clinical samples was determined by participating sites to be exempt from review or to not constitute human subjects research under applicable regulations, including 45 CFR §46.102. No samples were collected specifically for this study, and study findings were not used for patient care. The study was conducted in accordance with the principles of the Declaration of Helsinki.

## Use of AI-assisted Technologies

During the preparation of this work the authors used ChatGPT and CoPilot in order to assist with grammar and readability. After using this tool/service, the authors reviewed and edited the content as needed and take full responsibility for the content of the publication.

