## Supplemental Data for "Multisite Evaluation of an Amplification-based Nanopore Sequencing Solution to Analyze Challenging Clinically Relevant Variants in Genes Associated with Hereditary Diseases"

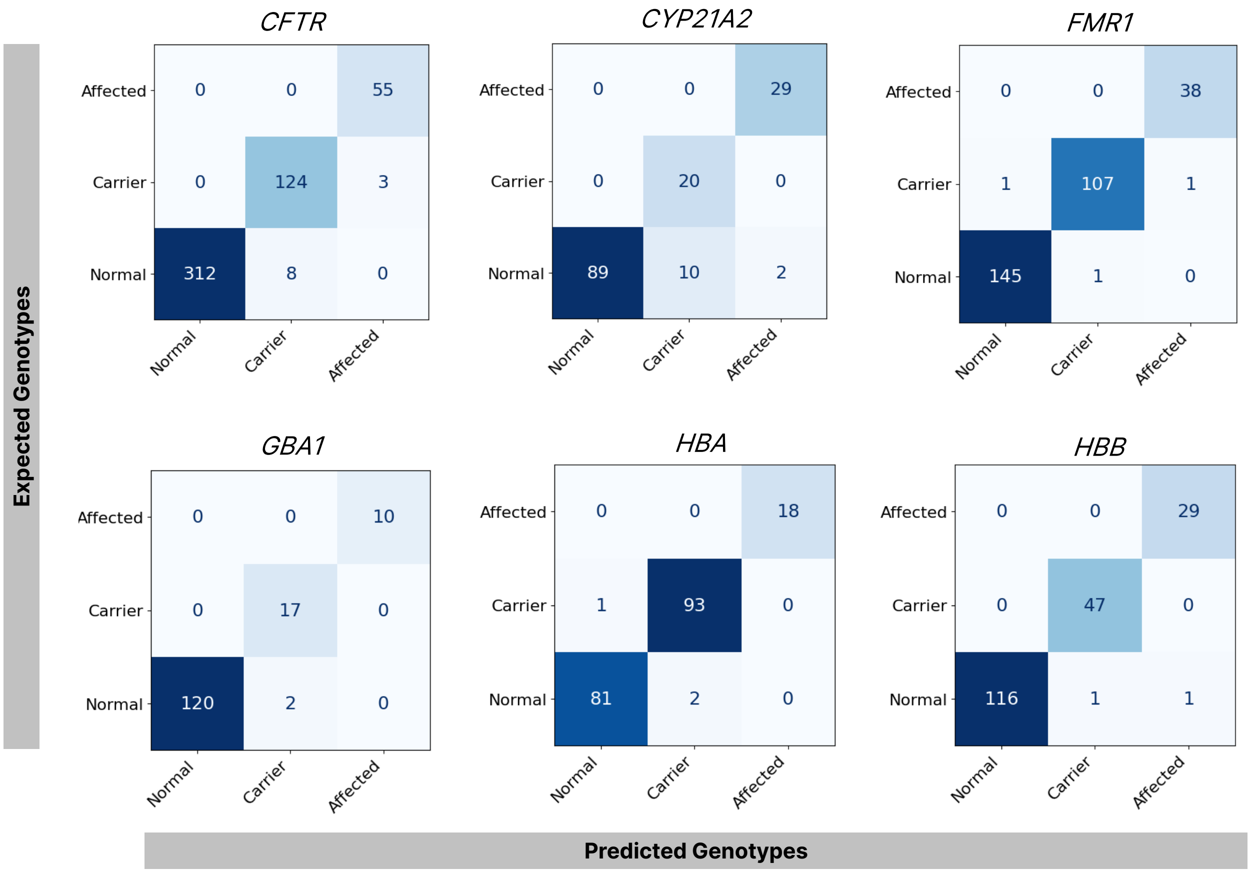


**Figure S1. No-exclusions genotype status contingency tables for genes with deviations from contingency tables described in Table 1.** Discrepant samples lacking gene-level orthogonal confirmation were excluded from the main text contingency tables (Figure 2) to avoid assumption-introduced bias, but are included above for 6/8 genes that diverge from main text.

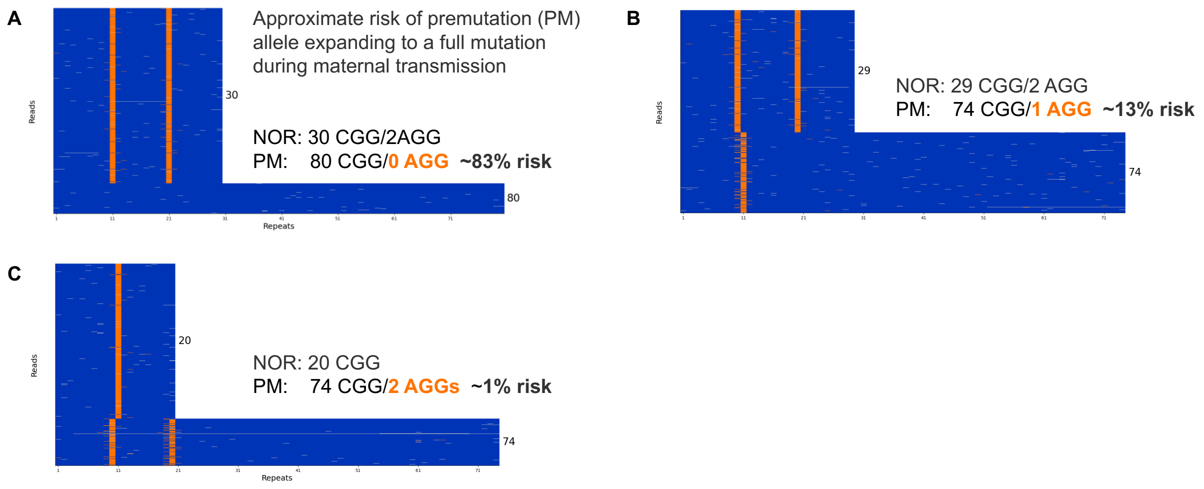


**Figure S2. Integrated CGG sizing and AGG phasing enable better expansion risk interpretation in *FMR1* premutation samples.** Simultaneous determination of CGG repeat length and phasing of AGG interruptions in *FMR1* premutation alleles provides clinically important information for carrier screening, as AGG interruptions reduce the likelihood of expansion to a full mutation during maternal transmission.[21-23] This information is most relevant for premutation alleles with fewer than 100 CGG repeats, with the greatest differences in predicted expansion risk observed in the 70–80 CGG range.

Shown are three representative female premutation samples from this study with similar CGG repeat lengths within this range but differing AGG interruption patterns. For each sample, a waterfall plot of long-read amplicons grouped by size (blue) with AGG interruptions highlighted (orange) enables visualization of AGG number and position within the CGG tract. Despite comparable CGG repeat lengths, predicted expansion risk varied substantially across these samples (1%–83%) depending on AGG interruption patterns. Results shown are concordant with orthogonal CGG sizing (AmplideX PCR/CE) followed by reflex AGG testing (AmplideX PCR/CE *FMR1* Xpansion Interpreter® testing service; Asuragen), and corresponding expansion risk estimates are annotated for each sample.

**A**

| Genotype | Rgn01 | Rgn02 | Rgn03 | Rgn04 | Rgn05 | *HBA2* (Rgn06) | Rgn07 | *HBA1* (Rgn08) | Rgn09 | Rgn10 | Rgn11 | Rgn12 | Rgn13 | Rgn14 |
|---|---|---|---|---|---|---|---|---|---|---|---|---|---|---|
| aa/aa | 2 | 2 | 2 | 2 | 2 | 2 | 2 | 2 | 2 | 2 | 2 | 2 | 2 | 2 |
| THAI/aa | 2 | 2 | 1 | 1 | 1 | 1 | 1 | 1 | 1 | 1 | 1 | 2 | 2 | 2 |
| MED-II/aa | 2 | 2 | 1 | 1 | 1 | 1 | 1 | 1 | 1 | 2 | 2 | 2 | 2 | 2 |
| FIL/aa | 2 | 2 | 1 | 1 | 1 | 1 | 1 | 1 | 1 | 1 | 2 | 2 | 2 | 2 |
| alpha20.5/aa | 2 | 2 | 1 | 1 | 1 | 1 | 1 | 1 | 2 | 2 | 2 | 2 | 2 | 2 |
| MED-I/aa | 2 | 2 | 2 | 1 | 1 | 1 | 1 | 1 | 1 | 2 | 2 | 2 | 2 | 2 |
| SEA/aa | 2 | 2 | 2 | 1 | 1 | 1 | 1 | 1 | 1 | 1 | 1 | 1 | 2 | 2 |
| 4.2del/aa | 2 | 2 | 2 | 2 | 1 | 1 | 2 | 2 | 2 | 2 | 2 | 2 | 2 | 2 |
| 3.7del/aa | 2 | 2 | 2 | 2 | 2 | 1 | 1 | 2 | 2 | 2 | 2 | 2 | 2 | 2 |
| Anti-4.2/aa | 2 | 2 | 2 | 2 | 3 | 3 | 2 | 2 | 2 | 2 | 2 | 2 | 2 | 2 |
| Anti-3.7/aa | 2 | 2 | 2 | 2 | 2 | 3 | 3 | 2 | 2 | 2 | 2 | 2 | 2 | 2 |
| 3.7del/anti-3.7 | 2 | 2 | 2 | 2 | 2 | 2 | 2 | 2 | 2 | 2 | 2 | 2 | 2 | 2 |
| 4.2del/anti-4.2 | 2 | 2 | 2 | 2 | 2 | 2 | 2 | 2 | 2 | 2 | 2 | 2 | 2 | 2 |
| 3.7del/anti-4.2 | 2 | 2 | 2 | 2 | 3 | 2 | 2 | 2 | 2 | 2 | 2 | 2 | 2 | 2 |
| 4.2del/anti-3.7 | 2 | 2 | 2 | 2 | 2 | 2 | 3 | 2 | 2 | 2 | 2 | 2 | 2 | 2 |
| Sample with rare 2-gene deletion | 2 | 2 | 2 | 2 | 2 | 1 | 1 | 1 | 1 | 1 | 1 | 1 | 2 | 2 |

**B**

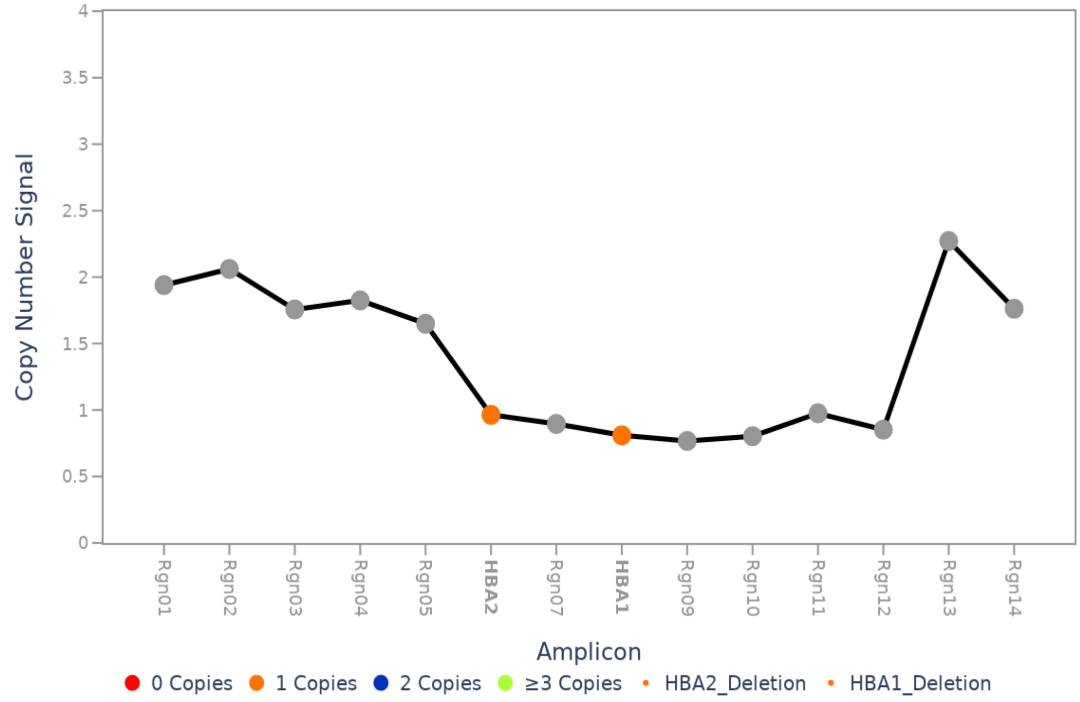


**Figure S3. Detection of α-globin cluster structural variants and copy-number profiles.** The Carrier Plus assay detects and classifies common α-globin cluster structural variants (SVs) and reports *HBA1* and *HBA2* copy number. **(A)** Normalized copy-number table across amplicons for samples with known α-globin genotypes, highlighting common SV patterns (blue and orange)

and a sample with a rare two-gene deletion profile (purple). **(B)** Software output showing copy-number signal profile for a sample with single-copy *HBA1* and *HBA2*; approximate breakpoint positions were inferred from transitions in amplicon copy-number signals.

**Table S1. Summary of orthogonal methods used for gene-specific variant confirmation across participating laboratories.** Methods span a broad range of molecular technologies, including Amplification-Refractory Mutation System PCR (ARMS), droplet digital PCR (ddPCR), capillary electrophoresis (CE), agarose gel electrophoresis (AGE), multiplex ligation-dependent probe amplification (MLPA), next-generation sequencing (NGS), long-range PCR, microarray-based approaches, quantitative PCR (qPCR), Sanger sequencing, strip assays, mass spectrometry, and whole-genome sequencing (WGS). In total, 16 distinct method types were represented across laboratories, reflecting site-specific testing workflows and the technical complexity of the genes evaluated. Product identifiers are listed where available and could be disclosed per laboratory policy.

| Gene | Method Type | Kit, Service, or Method Description | Product Identifier |
|---|---|---|---|
| *CFTR* | ARMS | Yourgene® Cystic Fibrosis Base | CF2EUB2 |
| *SMN1/2* | ddPCR | SMN1 Copy Number Determination Kit (BioRad, Hercules, California) | 1863500 |
| *SMN1/2* | ddPCR | SMN2 Copy Number Determination Kit (BioRad, Hercules, California) | 1863503 |
| *HBB* | GAP PCR | In-house | N/A |
| *HBA1/2* | GAP PCR/AGE | In-house based on PMID: 11439976 | N/A |
| *GBA* | Long-range PCR | Long-range PCR | N/A* |
| *CFTR* | Luminex | Luminex xTAG CF60 kit (Luminex Corp, Austin, Texas) | I024C0181 |
| *CFTR* | Microarray | CytoScan HD Microarray (Thermo Fisher Scientific, Waltham, Massachusetts) | 901835 |
| *CFTR* | Microarray | Illumina Infinium Bead chip array | GSAMD-24v2-0-Psych-24v1-1 |
| *CFTR* | MLPA | Multiplex Ligation Probe Amplification (MLPA) using MRC Holland SALSA probemix | N/A* |
| *CFTR* | MLPA | SALSA MLPA Probemix P091 CFTR (MRC Holland, Amsterdam, the Netherlands) | P091 |
| *SMN1/2* | MLPA | Multiplex Ligation Probe Amplification (MLPA) using MRC Holland SALSA probemix | N/A* |

| | | | |
|---|---|---|---|
| *SMN1/2* | MLPA | SALSA MLPA assays Probemix P021 and P460 (MRC Holland, Amsterdam, the Netherlands) | P021 and P460 |
| *HBA1/2* | MLPA | MRC Holland | P140_C1_HBA |
| *HBA1/2* | MLPA | Multiplex Ligation Probe Amplification (MLPA) using MRC Holland SALSA probemix | N/A* |
| *HBB* | MLPA | Multiplex Ligation Probe Amplification (MLPA) using MRC Holland SALSA probemix | N/A* |
| *HBB* | MLPA | SALSA MLPA assays Probemix P102 (MRC Holland, Amsterdam, the Netherlands) | P102 |
| *GBA* | MLPA | SALSA MLPA Probemix P338 GBA (MRC Holland, Amsterdam, the Netherlands) | P338 |
| *CFTR* | NGS | SureSelect Custom Target Enrichment for CFTR (Agilent Technologies) | custom |
| *CFTR* | NGS | TruSight Cystic Fibrosis 139-Variant Assay (Illumina Inc, San Diego, California) | 20036925 |
| *CFTR* | NGS | Illumina Custom-Designed Primary Ciliary Dyskinesia (PCD) NGS Test, including CFTR | custom |
| *CFTR* | NGS | Invitae Cystic Fibrosis Test | 04714 |
| *SMN1/2* | NGS | Next Generation Sequencing using Illumina | N/A* |
| *FMR1* | NGS | Next Generation Sequencing using Illumina | N/A* |
| *HBA1/2* | NGS | Next Generation Sequencing using Illumina | N/A* |
| *HBB* | NGS | Next Generation Sequencing using Illumina | N/A* |
| *CYP21A2* | NGS | Next Generation Sequencing using Illumina | N/A* |
| *TNXB* | NGS | Next Generation Sequencing using Illumina | N/A* |
| *GBA* | OpenArray | TaqMan OpenArray custom pannel | N/A* |
| *CFTR* | PCR/CE | AmplideX® PCR/CE CFTR Kit (Asuragen, a Bio-Techne Brand, Austin, Texas) | A00520 |

| | | | |
|---|---|---|---|
| *SMN1/2* | PCR/CE | AmplideX® PCR/CE SMN1/2 Plus Kit (Asuragen, a Bio-Techne Brand, Austin, Texas) | A00054 |
| *FMR1* | PCR/CE | AmplideX PCR/CE FMR1 Reagents (Asuragen, a Bio-Techne Brand, Austin, Texas) | 49402 |
| *FMR1* | PCR/CE | Xpansion Interpreter® Service (Asuragen, a Bio-Techne Brand, Austin, Texas) | CLIA lab service |
| *FMR1* | PCR/CE | PCR / capillary electrophoresis (CE) | N/A* |
| *F8* | PCR/AGE | Inverse PCR/gel electrophoresis | N/A* |
| *CFTR* | PCR/MALDI-TOF | Custom PCR followed by MALDI-TOF Mass Spectrometry using Aegena Bioscience MassARRAY | custom |
| *SMN1/2* | qPCR | Custom designed Taqman PCR (ThermoFisher Scientific, Waltham, Massachusetts) | custom |
| *CYP21A2* | qPCR | CAH RealFast™ CNV Assay (ViennaLab Diagnostics GmbH, Vienna, Austria) | 7-410 |
| *CFTR* | Sanger | Bidirectional Sanger sequencing | N/A |
| *SMN1/2* | Sanger | Bidirectional Sanger sequencing | N/A |
| *HBA1/2* | Sanger | Bidirectional Sanger sequencing | N/A |
| *HBA1/2* | Sanger | Bidirectional Sanger sequencing | N/A |
| *HBB* | Sanger | Bidirectional Sanger sequencing | N/A |
| *CYP21A2* | Sanger | Bidirectional Sanger sequencing | N/A |
| *GBA* | Sanger | Bidirectional Sanger sequencing | N/A |
| *HBB* | Strip assay (allele-specific hybridization assay) | Beta-Globin StripAssay® Kit IME (ViennaLab Diagnostics GmbH, Vienna, Austria) | 4-140 |
| *HBB* | Strip assay (allele-specific hybridization assay) | Beta-Globin StripAssay® Kit SEA (ViennaLab Diagnostics GmbH, Vienna, Austria) | 4-150 |

| | | | |
|---|---|---|---|
| *CYP21A2* | Strip assay (allele-specific hybridization assay) | CAH StripAssay® (ViennaLab Diagnostics GmbH, Vienna, Austria) | 7-380 |
| *CYP21A2* | WGS | PCR-Free whole genome seq, Illumina DRAGEN | N/A |

* Product identifier not disclosed per laboratory policy

**Table S2. CGG repeat sizing tolerance thresholds used in this study.** CGG repeat sizing tolerance thresholds applied in the evaluation of the AmplideX Nanopore Carrier Plus assay (Mix B) were intentionally more stringent than those defined by CAP/ACMG proficiency testing guidelines[23] and published recommendations to maintain consistency with tolerance criteria used in the orthogonal PCR/CE assays.

| Source | CGG Range | Tolerance |
|---|---|---|
| CAP/ACMG proficiency testing criteria;<br>ACMG Fragile X testing guidelines (2013);<br>Spector et al. (2021) | <55<br>56–100<br>>100 | ±5 repeats<br>±10 repeats<br>±2 SDs |
| AmplideX Nanopore Carrier Plus kit (Mix B) | <70<br>70-119<br>≥120 | ±1 repeat<br>±3 repeats<br>5% |

**Table S3. Logic for interpretation of software genotypes summary column to derive predicted genotype status for genes *CFTR*, *F8*, *FMR1*, *HBB*, and *SMN1*.** Acronyms and/or characters used to derive predicted genotype status ('Interpreted Genotype Status') from the 'Summary' column of the genotypes summary software analysis output ('Genotype Summary Acronym or Character'), based on biological interpretation and/or meaning ('Corresponding Abbreviation or Meaning') for applicable genes ('Applicable Genes'). 'Notes' column outlines gene-specific exceptions and associated handling.

| Genotype Summary Acronym or Character | Corresponding Abbreviation or Meaning | Interpreted Genotype Status | Applicable Genes | Notes |
|---|---|---|---|---|
| No Variants | No P/LP variants | Normal | *CFTR, F8, FMR1, HBB* | |
| V | Variant | Carrier | *CFTR, F8, HBB, SMN1* | |
| PM | Premutation | Carrier | *FMR1* | |
| 1 cp | 1 copy | Carrier | *SMN1* | |
| \| | Compound heterozygosity | Affected | *CFTR, F8, HBB, SMN1* | |
| , | Multiple variants | Affected | *CFTR, F8, HBB, SMN1* | Only copy number considered if 'LV' detected for *SMN1* |
| V(H) | Homozygous variant | Affected | *CFTR, F8, HBB, SMN1* | |
| 2V | 2 SNV/Indel | Affected | *CFTR, F8, HBB, SMN1* | |
| 2SV | 2 structural variants (fusions, inversions, large deletions, and insertions >50 bp) | Affected | *CFTR, F8, SMN1* | Assumed Carrier for *HBB* only |
| FM | Full mutation | Affected | *FMR1* | |
| 0 cp | 0 copies | Affected | *SMN1* | |

**Table S4. Variant-level agreement by gene and variant class across individual laboratories.** Site 1 represents the developer laboratory; Sites 2–5 represent external laboratories. Columns show OPA with orthogonal methods, followed by the number of measurements in parentheses. Variant types include SNVs/indels, copy numbers (CNs), large structural variants (SV), short tandem repeats (*FMR1* CGG and *CFTR* poly-T/TG) and interruptions (AGG), and category-level classifications (e.g., CGG category, alpha cluster CN category). OPA is equivalent to positive performance agreement (PPA) for select analyses (i.e., *CFTR* exon deletions and *F8* intron 1 and 22 inversions) due to the absence of orthogonal confirmation of negative samples. Abbreviations: V = variant-level concordance; S = sample-level concordance; A = allele-level concordance; I = interruption-level concordance; Exon Del = large exon deletion; PolyT/TG = poly-T/TG tract; Int1/22 Inv = intron 1/intron 22 inversion. NA = data not available or not applicable for a given site or variant type.

| Gene | Variant | | All 5 Sites | Site 1 | Site 2 | Site 3 | Site 4 | Site 5 |
|---|---|---|---|---|---|---|---|---|
| *CFTR* | SNV/Indel | (V) | 100.00% (34058) | 100.00% (20288) | 100.00% (316) | 99.91% (1086) | 100.00% (10726) | 100.00% (1642) |
| | LED | (S) | 100.00% (24) | 100.00% (10) | 100.00% (5) | 100.00% (4) | NA NA | 100.00% (5) |
| | PolyT/TG | (S) | 99.60% (278/224) | 100.00% (183) | 100.00% (2) | 97.50% (9) | 100.00% (20/18) | 98.03% (64/12) |
| *SMN1/2* | SNV/Indel | (V) | 99.80% (1515) | 100.00% (1009) | 100.00% (12) | 100.00% (85) | 100.00% (186) | 98.65% (223) |
| | *SMN1* CN | (S) | 98.25% (456) | 96.24% (186) | 97.83% (46) | 100.00% (62) | 100.00% (113) | 100.00% (49) |
| | *SMN2* CN | (S) | 98.25% (456) | 96.24% (186) | 100.00% (46) | 98.39% (62) | 100.00% (113) | 100.00% (49) |
| *FMR1* | CGG | (A) | 96.86% (541) | 98.33% (180) | 94.74% (57) | 96.58% (117) | 96.26% (187) | |
| | AGG | (I) | 100.00% (387) | 100.00% (222) | NA NA | 100.00% (144) | 100.00% (21) | |
| | CGG Category | (S) | 99.66% (292) | 100.00% (96) | 96.97% (33) | 100.00% (59) | 100.00% (104) | |
| *HBA1/2* | SNV/Indel | (V) | 100.00% (52) | 100.00% (19) | 100.00% (9) | 100.00% (24) | | |
| | *HBA1/2* CN | (S) | 99.32% (147) | 100.00% (66) | 97.50% (40) | 100.00% (41) | | |

| | | | | | | |
|---|---|---|---|---|---|---|
| | Alpha Cluster CN Category | (S) | 99.25% (134) | 100.00% (66) | 97.30% (37) | 100.00% (31) |
| *HBB* | SNV/Indel | (V) | 99.20% (125) | 99.03% (103) | 100.00% (22) | NA NA |
| | LED | (S) | 100.00% (48/46) | 100.00% (28) | 100.00% (20/21) | NA NA |
| *CYP21A2* | SNV/Indel | (V) | 100.00% (440) | 100.00% (426) | 100.00% (14) | |
| *SMN1/2* | SNV | (V) | 99.63% (267) | 99.63% (267) | NA NA | |
| *GBA1* | SNV/Indel | (V) | 100.00% (238) | 100.00% (220) | 100.00% (18) | |
| *F8* | Int1/22 Inv | (S) | 100.00% (11) | 100.00% (4) | 100.00% (7) | |

**Table S5. Variant calls for mix or mix combination of 136 cell lines available from Coriell Cell Repository.**

Data attached
